# The RUMIGEN EpiChip: a versatile, medium density DNA methylation Beadchip for large scale population studies in cattle

**DOI:** 10.64898/2025.12.19.695474

**Authors:** Valentin Costes, Adrian Lopez-Catalina, Amrita Raja Ravi Shankar, Gabriel Costa Monteiro Moreira, Sophie Martel, Ludivine Liétar, Carmen Garcia Alba, Javier L. Viana, Aurélie Chaulot-Talmon, Francesca Ali, Christine Couldrey, Mekki Boussaha, Sébastien Fritz, Eveline M. Ibeagha-Awemu, Nathalie Bissonnette, Jess Powell, Pau Navarro, Alex Caulton, Kumiko Takeda, Eiji Kobayashi, Gilles Foucras, Daniel Rico, Chrystelle Le Danvic, Laurent Schibler, Zexi Cai, Goutam Sahana, Shannon Clarke, Hélène Jammes, Oscar Gonzalez-Recio, Clotilde Patry, Hélène Kiefer

## Abstract

**Background:** DNA methylation contributes to the elaboration of phenotypes and is hypothesized to account for interindividual variations in farm animals. Currently, methodologies available to investigate DNA methylation in cattle rely on high throughput sequencing, which cannot be applied to large cohorts. To enable large scale DNA methylation analysis, we developed the RUMIGEN EpiChip, the first DNA methylation array specifically designed for cattle.

**Results:** Manufactured by Illumina, the EpiChip contains 43,317 CpG markers allowing analysis of phenotypes of agronomical interest as well as the study of regulatory processes. The assay design drew on data from numerous studies achieved by the scientific community, and includes CpGs where methylation varies with health status, physiological stage, fertility, and environmental challenges, as well as CpGs located in functional genomic elements (promoters, CTCF binding sites, expression quantitative trait loci). The technical performances of the EpiChip were tested on several semen and blood DNA samples and in two laboratories, showing excellent repeatability, accuracy and interoperability. Methylation values were also concordant with those obtained by reduced representation bisulfite sequencing. The EpiChip has demonstrated a good ability to explore biological processes such as genomic imprinting and differences between cell types, opening the possibility of inferring blood cell composition. Finally, analysis of longitudinal ear biopsies allowed accurate age prediction, suggesting the potential of the array to refine epigenetic clocks.

**Conclusions:** The RUMIGEN EpiChip is a cost-effective and versatile resource that opens new opportunities for the study of regulatory mechanisms underlying phenotypic variation and for large-scale association analyses in cattle.

## Background

DNA methylation, the most extensively studied epigenetic modification, occurs primarily at cytosine followed by guanine (CpG) sites. The genomic distribution, density, and methylation status of CpGs influence chromatin architecture and regulate gene expression. DNA methylation plays critical roles in cell fate determination and development, chromosome structure maintenance and repression of transposable elements. Significant deviations in DNA methylation landscapes can alter cell identity and compromise genome stability, leading to developmental disorders, infertility or cancers (1–3). Beyond these fundamental roles, DNA methylation is also influenced by environmental factors, aging, and physiological condition; however, such changes are typically modest in magnitude and affect only a limited subset of CpGs across the genome. Detecting these changes requires highly sensitive, quantitative, genome-wide methods that are cost-effective enough to be applied to relatively large sample sizes, ensuring adequate statistical power.

Whole genome bisulfite sequencing (WGBS), the gold standard method for genome-wide DNA methylation analysis, involves the conversion of unmethylated cytosines to thymines using sodium bisulfite followed by short-read sequencing (4–6). A more recent method that uses enzymatic conversion offers an alternative to sodium bisulfite treatment (7). However, both approaches require sequence alignment to an *in silico*–converted reference genome—a process complicated by the near-complete loss of cytosines after conversion, reflecting the low methylcytosine content of mammalian genomes. An additional drawback of these methods is that C/T single nucleotide polymorphisms (SNPs) can be misinterpreted as conversion events. Nanopore sequencing overcomes these limitations by enabling the direct detection of methylated cytosines (8) and, additionally, producing long reads that facilitate the analysis of repetitive elements (9). Across all these technologies, methylation at each CpG is quantified based on the proportion of reads identified as “methylated”, which depends on the total number of reads (“methylated” plus “unmethylated”) covering that site. The requirements for adequate sequencing depth, coupled with maximal genome coverage, contribute to the high cost and substantial computational demands, therefore limiting scalability of these techniques to large sample sizes. Alternative approaches have been developed to focus sequencing on restricted genomic regions. These include reduced representation bisulfite sequencing (RRBS), which preferentially targets CpG-dense features (10), and capture-based assays, which allow customization of the targeted regions (11).

DNA methylation arrays that allow for the analysis of thousands of CpG sites in a large number of samples have evolved alongside sequencing-based approaches. Historically, the development of methylation BeadChips began with the Human 1500 CpG GoldenGate assay, quickly followed by the release of the 27K BeadChip (12, 13). The design of these early arrays focused primarily on gene promoters and selected cancer-related loci. Few years later, a new version of the array was released, targeting about 450K CpGs (14). This updated design increased genome coverage by enhancing probe density in gene promoters and gene bodies, and including intergenic CpGs. The most recent version of the human DNA methylation BeadChip, the MethylationEPIC v2.0 array, includes approximately 935,000 CpG sites (15). Its design significantly expanded the coverage of intergenic CpG sites where distal regulatory elements such as enhancers are frequently found. The BeadChip methylation arrays have been widely used in large-scale human studies for over a decade. These assays capture sodium bisulfite–converted DNA using probes terminating at a CpG site. Hybridization is followed by a single-base extension step with fluorescence incorporation, the outcome of which reflects the methylation status of the targeted CpG (16). While less comprehensive in coverage than whole-genome sequencing methods, Infinium array–based methods remain the preferred tool for epigenome-wide association studies (EWAS) in humans (17). They are high-throughput, relatively affordable, and compatible with SNP genotyping platforms (18). Their long-standing use has revealed unprecedented subtle DNA methylation changes associated with complex traits, disease onset, and adverse environmental exposures in humans (19).

Cattle are a livestock species of major agroeconomic importance. Over recent decades, conventional selection, followed by genomic selection based on SNP array genotyping, has led to substantial improvements in performance across a wide range of traits (20). More recently, epigenetic mechanisms have emerged as a potential source of phenotypic variance in farm animals, that is not captured by genomic data alone (21). A variety of sequencing-based approaches have been employed in cattle to identify DNA methylation changes associated with diverse phenotypes and environmental factors, including mastitis and milk production (22–25), early-life nutrition and fertility in males (26–29), stress induced by maternal lactation (30), gut immunity (31) and immune cell differentiation (32), among others. Despite these advances, translating these epigenetic insights into practical applications for the improvement of management or breeding remains challenging. To address this, a high-throughput, robust, and cost-effective assay is urgently needed to investigate, in sufficiently powered studies, the association between DNA methylation, phenotypic and environmental variations, and ultimately to integrate methylation data into management and breeding strategies (33–35).

In this study, we introduce the RUMIGEN EpiChip, the first medium-density DNA methylation array developed for cattle. We describe its design and genome coverage and assess its technical performance. We also demonstrate its utility for investigating several biological processes in which DNA methylation is involved, including genomic imprinting, cell type-specific differences, and aging.

## Results

### Selection of the CpG content

Array-based technologies have the limitation of interrogating a predetermined set of CpGs. To develop a cost-effective and versatile platform, a format comprising 50,000 probes was selected, targeting CpGs relevant for both field applications and the investigation of biological questions. Two complementary selection strategies were implemented. The first focused on selecting CpG biomarkers linked to agronomically relevant traits, where DNA methylation variation associated with phenotypic and environmental changes was previously detected. To complement this first subset, the second strategy aimed to select CpGs located within key genomic features, thus enhancing the potential of the EpiChip to explore the role of DNA methylation in regulatory processes (Figure 1).

**Figure 1.**
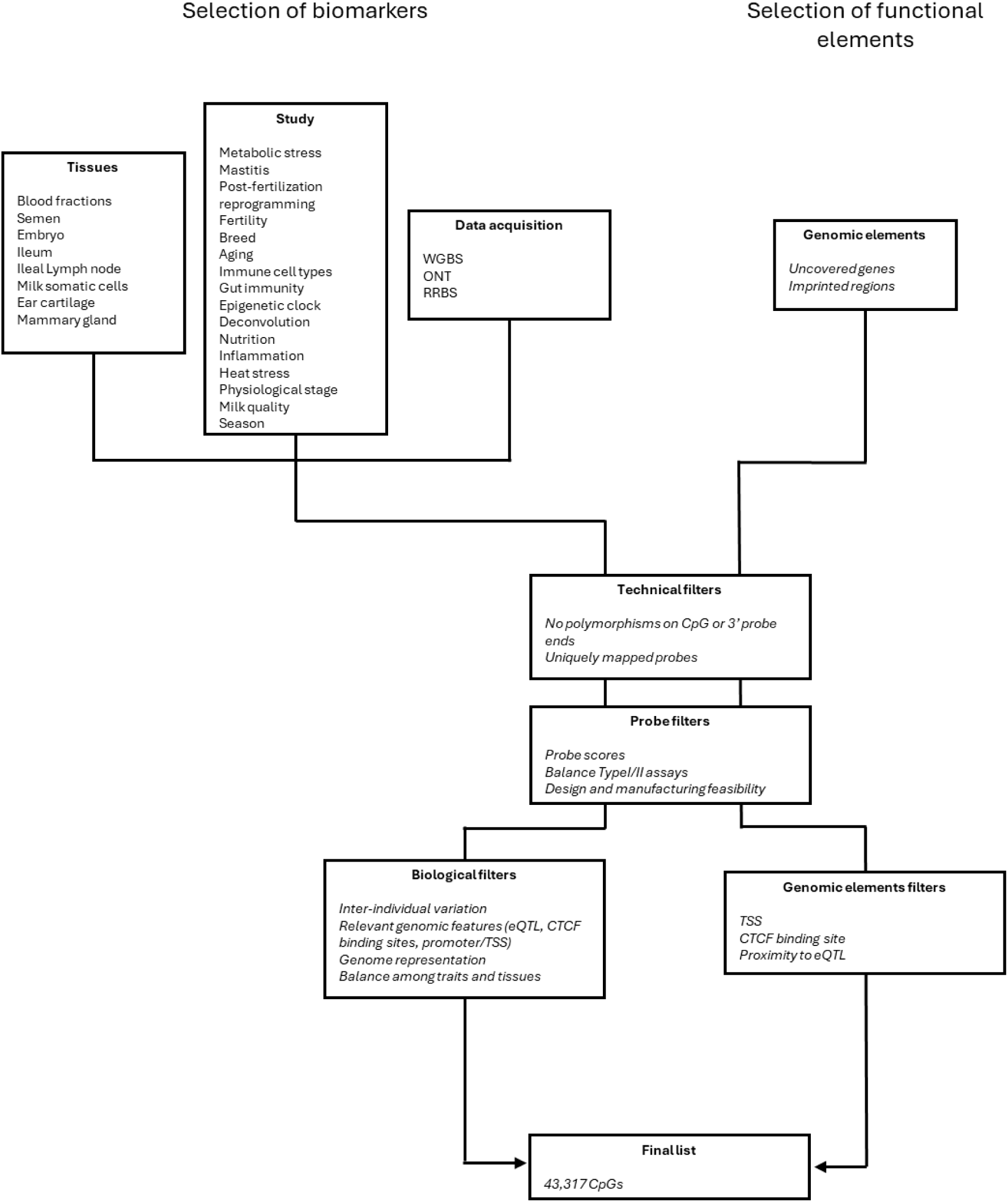
Workflow of the RUMIGEN EpiChip design strategy. Type I assays use two beads per probe for each targeted CpG (one probe for the methylated allele in the red channel and one probe for the unmethylated allele in the green channel). Type I assays use one bead per probe and per CpG (in either the red or green channel representing the methylated or unmethylated allele, respectively). RRBS, reduced representation bisulfite sequencing; ONT, Oxford Nanopore Technology; WGBS, whole genome bisulfite sequencing; TSS, transcription start site; eQTL, expression quantitative trait locus; CTCF, CCCTC-binding factor.

To support a biomarker-driven strategy, CpG sites associated with health, physiological stages, fertility and environmental challenges were compiled. These CpGs were identified by integrating data generated from multiple platforms— including RRBS, Nanopore sequencing, mammalian methylation array and WGBS—across a diverse set of tissues such as blood, semen, ileum, ileal lymph node, mammary gland, milk somatic cells, ear biopsies and the embryo, leading to a collection of 250K unique CpG sites that were subsequently used for probe design (Supplementary Table 1).

After the design, approximately 30K CpGs were discarded as no appropriate probes could be designed, but the number of designable probes was still too numerous for a medium density BeadChip design targeting approximately 50k probes. To reduce the list, three types of filters were applied, technical filters, probe filters and biological filters. Technical filters removed probes with SNPs within the CpG sites or within the last 15 nucleotides, as well as probes that mapped to several locations on an *in-silico* bisulfite-converted genome, which both constitute known biases that could introduce erroneous signals (15). Probe filters incorporated probe scores—intended to predict the likelihood of generating high-quality signals—and the balance between Type I and Type II assays, which is important for background normalization during preprocessing. Finally, biological filters relied on available data and biological annotations to select the most relevant CpGs among all those that passed the previous filters. CpGs were selected based on high inter-individual variability, location within regulatory elements (e.g., promoters, transcription start sites) and proximity to expression quantitative trait loci (eQTLs) (36) and CCCTC-binding factor (CTCF) binding sites (37). In addition, to reduce redundancy from closely spaced CpGs, only one representative was retained within a recursive 10-bp window. This approach ensures spatial uniqueness and promotes a more balanced genomic coverage. To achieve an inclusive representation of traits, tissues, and genome regions, the stringency of the biological filter was adjusted taking account of the CpG subsets listed in Supplementary Table 1. The origin of the selected biomarkers in terms of identification techniques, traits and tissues is shown in Supplementary Figure 1.

To preserve the ability to explore biological processes beyond those targeted in the biomarker-based approach, a second strategy was adopted to improve the genome-wide representativeness of the array. This annotation-based strategy targeted CpGs within uncovered gene promoters and imprinted regions without prior evidence of any association with phenotypes. After probe design, this approach used the same filters as the biomarker-based strategy but with slight modifications in the biological filters: spatial uniqueness and proximity to regulatory elements (eQTLs and CTCF binding sites) were considered, whereas interindividual variability was not, due to the lack of data for these regions; in addition, only one CpG per gene promoter was retained.

Upon completion of these design steps and manufacturing, the RUMIGEN EpiChip included a total of 44,311 markers, among which are 43,317 CpGs of the bovine genome, and a total of 11.6% of Type I probes.

### Genome coverage and regulatory features

The RUMIGEN EpiChip roughly covers all chromosomic regions (Supplementary Table 2 and 3). We then assessed the overlap between CpG sites present on the EpiChip and different genomic annotations by evaluating whether the observed CpG–feature associations are biologically specific or explainable by random CpG positioning.

In the analysis of commonly annotated features, we observed an enrichment (observed odds ratio; OR > 1) in promoters, transcription star sites (TSS_100bp), and transcription termination sites (TTS_100bp). These enrichments are specific, as the observed ORs lie clearly outside the confidence intervals derived from random CpG sets (Figure 2 and Supplementary Table 2). Conversely, exonic and intronic regions showed significant depletion, which was also specific according to the same randomization-based criterion. Notably, when enrichment was assessed using novel annotation catalogs, we observed enrichment for Enhancer_CAGE, Promoter_Ensembl, and open chromatin regions, together with depletion for TSS_CAGE, Enhancer_Ensembl, and ATAC regions (Figure 2 and Supplementary Table 3). However, the enrichment, although present, was not specific, as comparable signals were obtained from randomly generated CpG sets. Thus, the CpGs included on the EpiChip do not show biologically specific enrichment and largely mirror the genome-wide distribution of these novel annotations.

**Figure 2.**
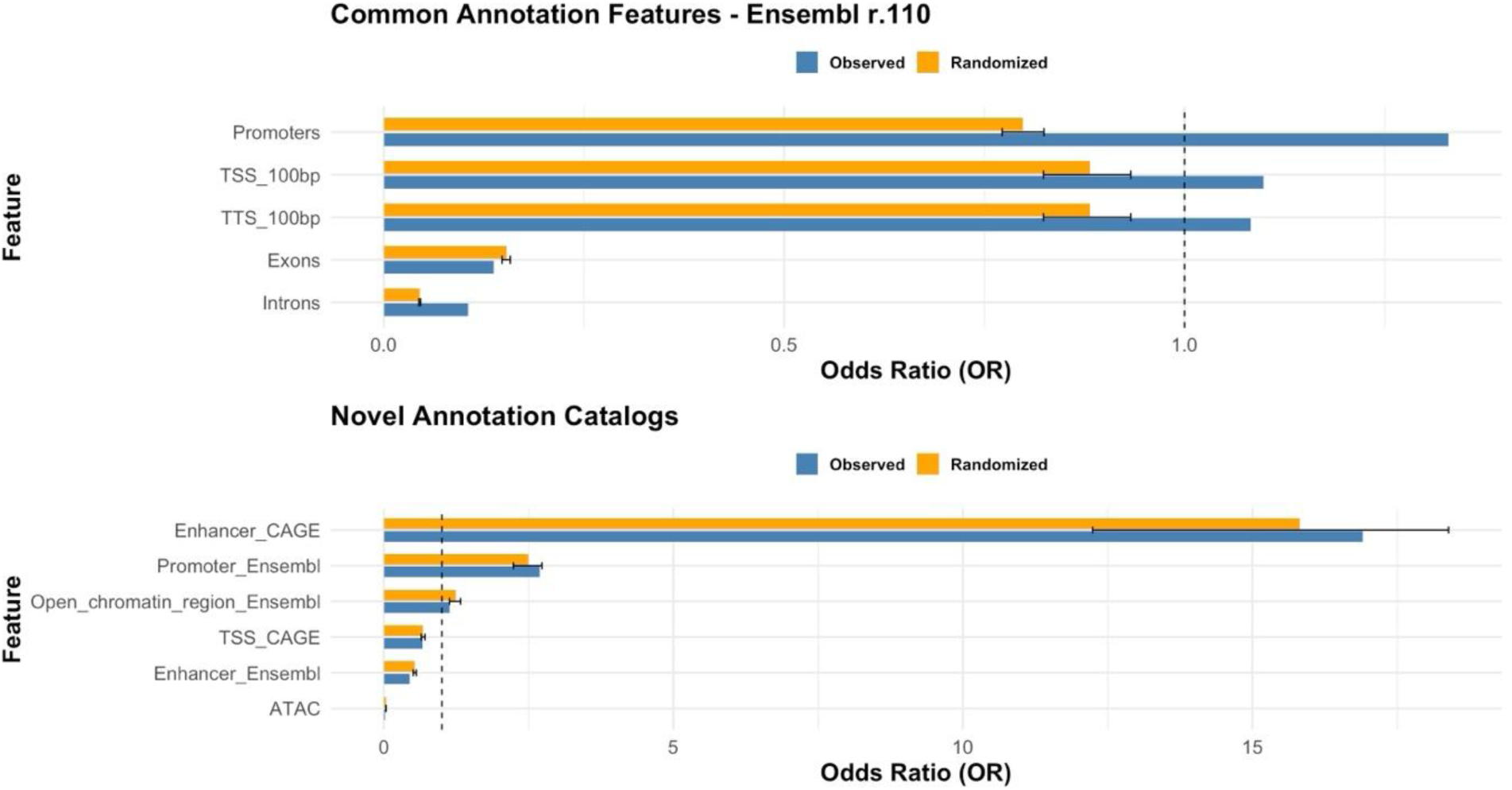
Enrichment of CpG sites in common and novel genomic annotations. The figure shows odds ratios (ORs) for overlap between the RUMIGEN EpiChip CpG sites and annotated genomic features. The top panel corresponds to commonly annotated features from Ensembl (release 110), and the bottom panel corresponds to novel annotation catalogs. For each feature, blue bars represent the observed OR, and orange bars represent the mean OR obtained from 1,000 randomized permutations of the same-sized background regions. Black horizontal lines on the orange bars indicate the 95% confidence intervals of the randomized ORs. The dashed vertical line at OR = 1 indicates the expected overlap under the null hypothesis. TSS, transcription start site; TTS, transcription termination site; CAGE, cap analysis of gene expression; ATAC, assay for transposase-accessible chromatin.

### Repeatability

To assess the array’s technical performance, we selected two easily accessible cell types that allow non-invasive sampling and provide sufficient genomic DNA for reliable technical replication: semen and blood cells.

An important aspect to evaluate is the repeatability of methylation measurements across technical replicates. To this end, two individual samples per cell type—semen samples #1 and #2 and blood samples #1 and #2—were analyzed, with 24 and 12 technical replicates per sample, for blood and semen, respectively. Within each cell type, correlations among the technical replicates were calculated and the results are presented in Supplementary Figure 2. The mean correlation was 0.993 for blood sample #1, 0.996 for blood sample #2 and 0.997 for both semen sample #1 and semen sample #2. These high correlation values across samples within both tissue types demonstrate the high reproducibility of the methylation measurements on the same DNA matrices.

### Concordance with reduced representation bisulfite sequencing

Although repeatability is very high, technical biases could arise and limit direct comparisons of methylation results obtained by the RUMIGEN EpiChip with those from other technologies. To address this, EpiChip DNA methylation measures (beta values, a numerical measure of DNA methylation, ranging from 0 to 1, calculated as the ratio of methylated signal intensity to the total intensity of both methylated and unmethylated signals) were compared to the methylation percentages obtained using bisulfite sequencing-based technology. For this analysis, eight samples were analyzed with RRBS and replicated across the EpiChip. Correlations between the RRBS data and EpiChip beta values were calculated for 11,207 to 15,412 CpGs analyzed with both technologies, with results presented in Figure 3A. The correlations are consistently high (average Pearson correlation > 0.89), indicating strong cross-technology reproducibility. Variations between the two methods may stem from the inherent challenges of obtaining precise methylation estimates with sequencing-based techniques such as RRBS, due to the limited sequencing depth at each CpG. As illustrated by the correlation plots obtained for two representative blood and semen samples, the strong correlations seem largely driven by hypomethylated and hypermethylated CpGs (Figure 3B, C). We therefore focused on intermediary methylation values (CpGs with methylation percentages between 20% and 80%). The correlations were still relatively high (R=0.73 for semen sample and 0.68 for blood sample; Figure 3D, E), further reinforcing confidence in the EpiChip.

**Figure 3.**
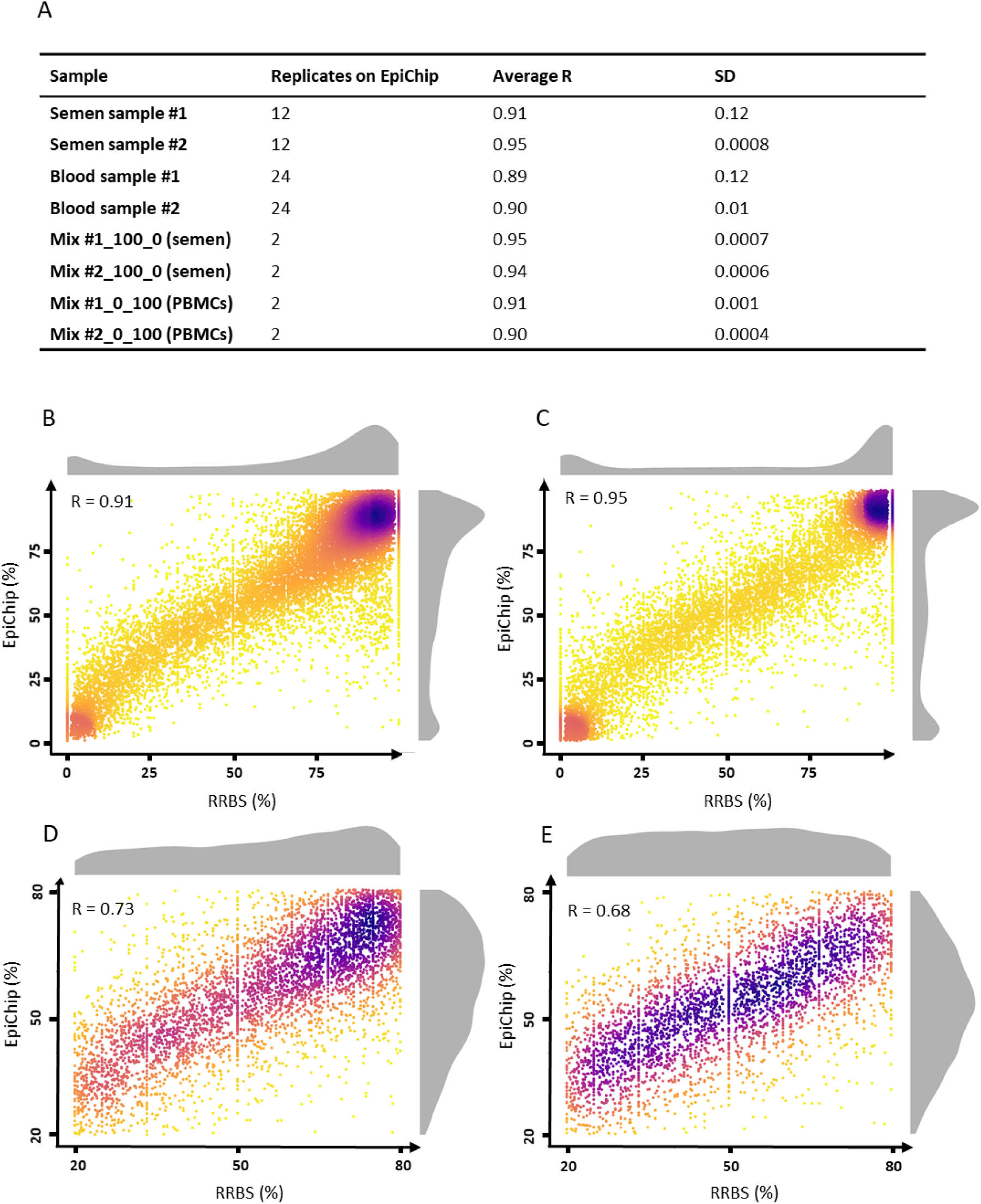
Concordance of the RUMIGEN EpiChip data with reduced representation bisulfite sequencing data. A: Correlation between the DNA methylation values of 8 samples measured by the EpiChip and by RRBS. Refer to Table 1 (Methods) for sample names. As samples have been analyzed several times with the EpiChip, the number of replicates, average Pearson’s correlation coefficient R and standard deviation (SD) are reported. B, C: Pairwise correlation plots between the DNA methylation percentages obtained with each method for blood sample #1 (B) and semen sample #2 (C). D, E: Pairwise correlation plots between the DNA methylation percentages obtained with each method at intermediary methylated CpGs for blood sample #1 (D) and semen sample #2 (E). Beta values are converted into methylation percentages. Pearson’s correlation coefficients are indicated and the distribution of methylation values obtained with each method is displayed in grey.

### Accuracy of the DNA methylation measures

Another key aspect evaluated was the accuracy of the array’s measurements, i.e. the ability to accurately detect known DNA methylation levels. To test this, we took advantage of paired blood and semen samples from two bulls. Pure DNA from semen and blood was used to prepare mixtures with blood DNA proportions ranging from 0% to 100%, in 10% increments, with the remaining portion consisting of semen DNA. Both pure and mixed DNA samples were analyzed with the EpiChip. Based on the known methylation levels in the pure samples and the dilution ratios in the mixtures, theoretical methylation levels for each mixture were calculated. The correlation between these theoretical methylation levels and the measured levels was very high (Pearson’s correlation coefficient R > 0.984; Figure 4A), and the median difference between theoretical and measured levels was 2.3%, with only 20% of CpGs displaying more than 5% difference between expected and measured methylation values (Figure 4B). This finding demonstrates that the RUMIGEN EpiChip has strong accuracy, enabling precise estimation of methylation levels at targeted CpG sites.

**Figure 4.**
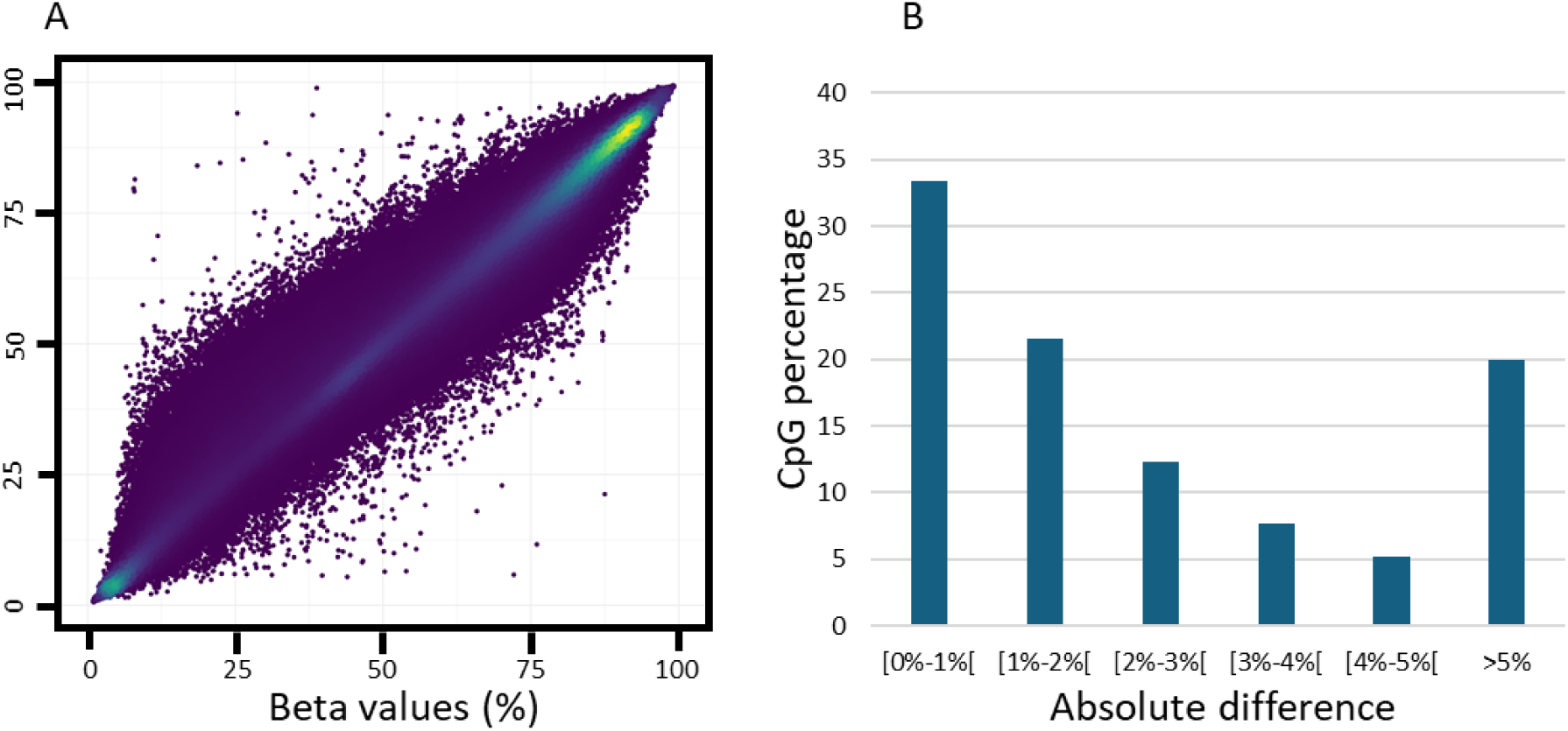
Accuracy of DNA methylation measures with the RUMIGEN EpiChip. Pure semen and blood samples from two individual bulls were analyzed using the RUMIGEN EpiChip. Next, mixed DNA samples were prepared by mixing 90%-10% of semen DNA with 10%-90% of blood DNA, with a 10% increment. Based on the methylation values on pure samples, and the proportion of each DNA in the mix, an expected methylation value was calculated. The mixed DNA samples were analyzed with the RUMIGEN EpiChip, and the beta values obtained were compared with the expected methylation values (n=1,444,212 values compared in total). A: pairwise correlation plot between the expected methylation values and the measured beta values. The Pearson’s correlation coefficient is above 0.98. B: Absolute differences were calculated between the expected methylation values and measured beta values (x-axis), and the CpG percentages found within each category is displayed (y-axis).

### Interoperability between batches and laboratories

Another key question was whether we could distinguish variance attributable to biological factors from that due to technical factors in a given laboratory and across laboratories. To investigate this, the same semen and blood samples as above were processed by two different laboratories, with two batches per laboratory, to examine variability arising from technical effects. A principal component analysis (PCA) was performed, and the first four dimensions are shown in Figure 5. As expected, the first dimension reflects differences between semen and blood. The second dimension shows that a portion of the variance is due to inter-laboratory differences, and to a lesser degree to the batch (batch effect visible only for Laboratory 1). Differences between the two blood samples appear on dimension 3, while differences between the two semen samples appear on dimension 4. This analysis highlights the presence of biological variability among both cell types and individuals, along with technical variation primarily due to inter-laboratory differences.

**Figure 5.**
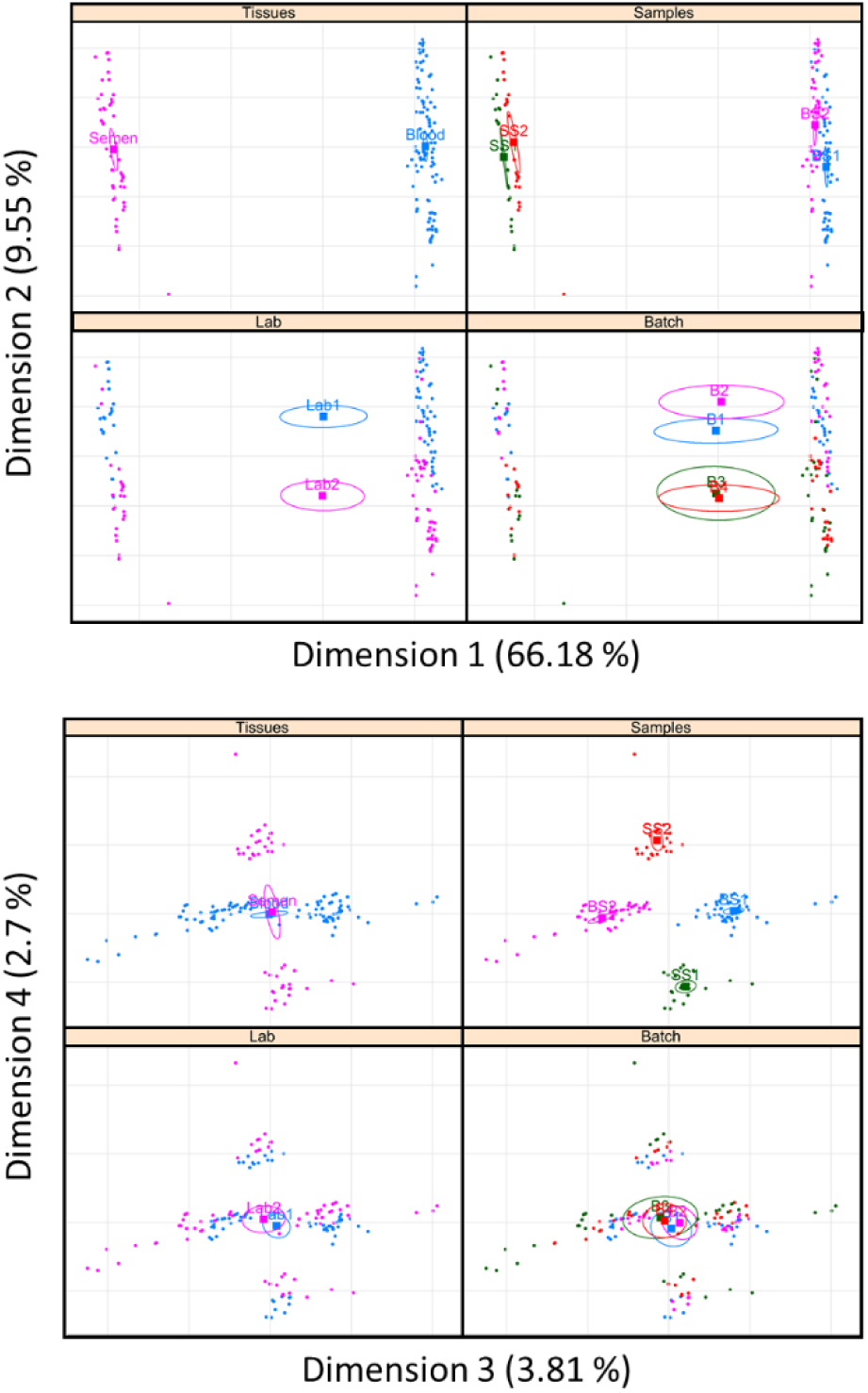
Interoperability of the RUMIGEN EpiChip. For two animals, two different cell types (semen and blood) were analyzed using the RUMIGEN EpiChip, resulting in four independent samples: BS1 and BS2-blood samples #1 and #2; SS1 and SS2-semen samples #1 and #2. For blood and semen, 48 and 24 technical replicates were analyzed per animal; respectively. These replicates were processed by two different laboratories in two different batches. A principal component analysis (PCA) was conducted. The individual factor maps are shown for the first four dimensions of the PCA, with the individuals displayed in different colors according to the cell type (upper left panel), sample (upper right panel), laboratory (lower left panel) and batch (lower right panel).

### Genomic imprinting

The RUMIGEN EpiChip was next used to investigate several regulatory processes in which DNA methylation is involved. We first focused on genomic imprinting, a phenomenon that leads to monoallelic, parent-of-origin expression of genes essential for development and growth, with DNA methylation playing a central role in the establishment and maintenance of imprinted loci.

The methylation status of blood and semen was investigated at 37 CpGs located in specific regions of 8 imprinted genes (Supplementary Table 4). For all regions, methylation in blood reached the expected value of 50%, consistent with the presence of two parent-of-origin alleles, one fully methylated and the second fully unmethylated. In contrast, in semen, the paternally expressed genes *GNAS, IGF2R, PEG3, PEG10, PLAGL1* and *SNRPN* showed weak methylation whereas the *IGF2/H19* imprinting control region (ICR) was fully methylated (Figure 6A). Methylation of the paternal allele at this ICR prevents the binding of the insulator CTCF and blocks the expression of paternal *H19*, allowing the activation of paternal *IGF2* from downstream enhancers (38). The sizes of the boxplots in Figure 6A reflect a certain degree of variability across the individual CpGs analyzed in each imprinted region (Figure 6B and Supplementary Figure 3), whereas at each single CpG the methylation levels across biological and technical replicates are very stable. These results suggest that the RUMIGEN EpiChip could be confidently used for investigating epigenetic plasticity at imprinted regions.

**Figure 6.**
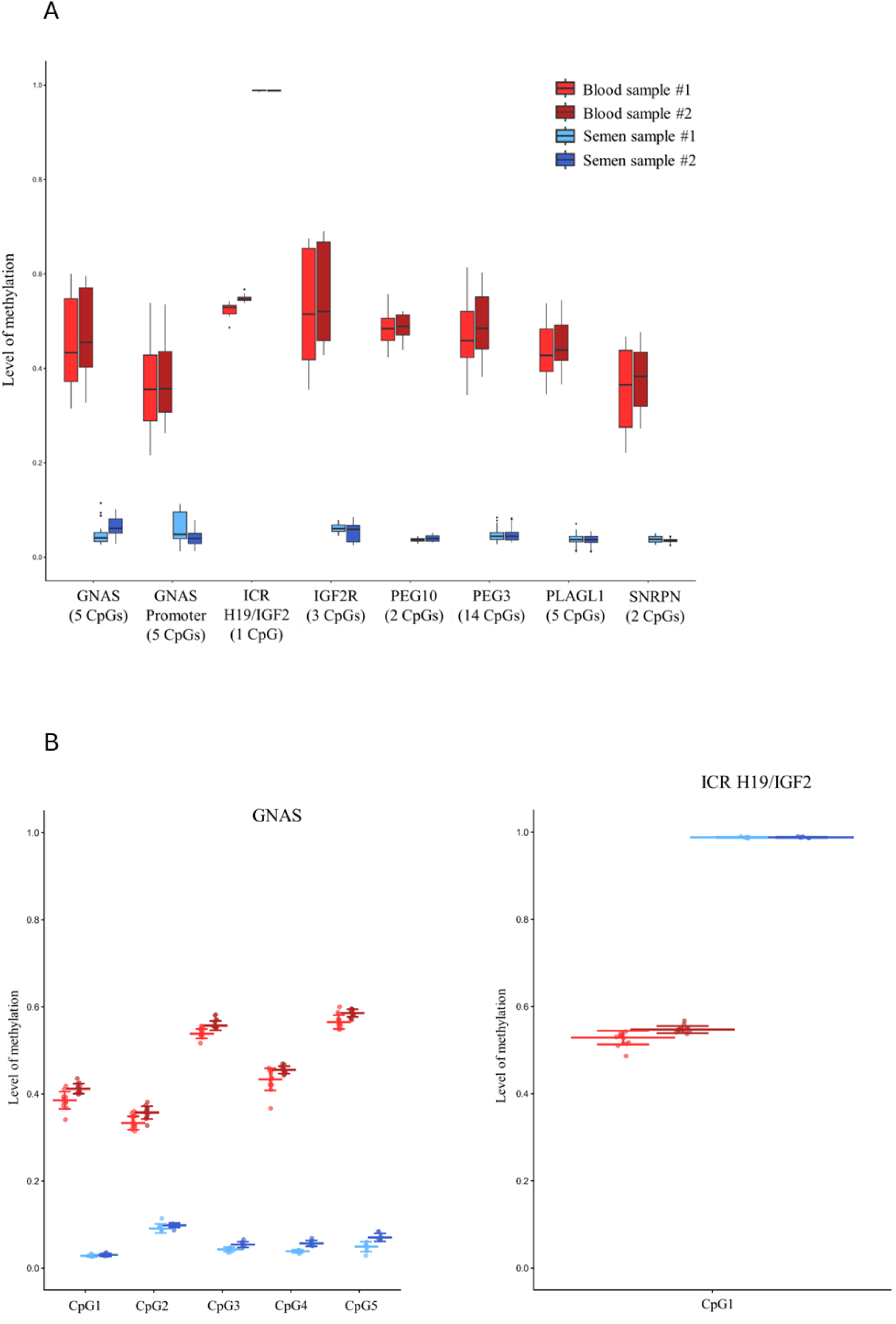
DNA methylation at imprinted regions in 8 genes analyzed using the RUMIGEN EpiChip. Technical replicates of four independent samples (blood samples #1 and #2 and semen samples #1 and #2, Table 1) from two different cell types (semen and blood) and two animals, were analyzed by one laboratory in one single batch. A: Mean DNA methylation values across 1 to 14 CpGs included in the imprinted regions. The number of CpGs analyzed is indicated in brackets for each region. B: For two regions located in *GNAS* and *IGF2R*, DNA methylation values are displayed at the level of single CpGs, revealing heterogeneity across CpGs but homogeneity within biological and technical replicates. Other regions are displayed in Supplementary Figure 3.

### Cell-type specific DNA methylation patterns

DNA methylation plays a key role in regulating transcription in a cell type-specific manner. As a consequence, the sperm methylome differs markedly from that of somatic cells, reflecting the specificity of the germline. We compared semen samples with blood samples and detected DNA methylation differences in germline genes (Supplementary Figure 4), consistent with earlier studies using sequencing-based approaches (39, 40). In the same line, blood is composed of several nucleated cell populations, each displaying its own methylation pattern. We compared the DNA methylation levels of neutrophils, monocytes and lymphocytes lineages, on a reference panel of CpGs able to discriminate each cell type and selected to conduct deconvolution approaches (Costes et al., personal communication; under reference DI-RV-24-0038, INRAE’s Intellectual Property and Valorization Committee). The DNA methylation patterns at these CpGs are very distinct between cell types (Figure 7), suggesting that deconvolution approaches can be applied to EpiChip data obtained on whole blood to infer cell composition.

**Figure 7.**
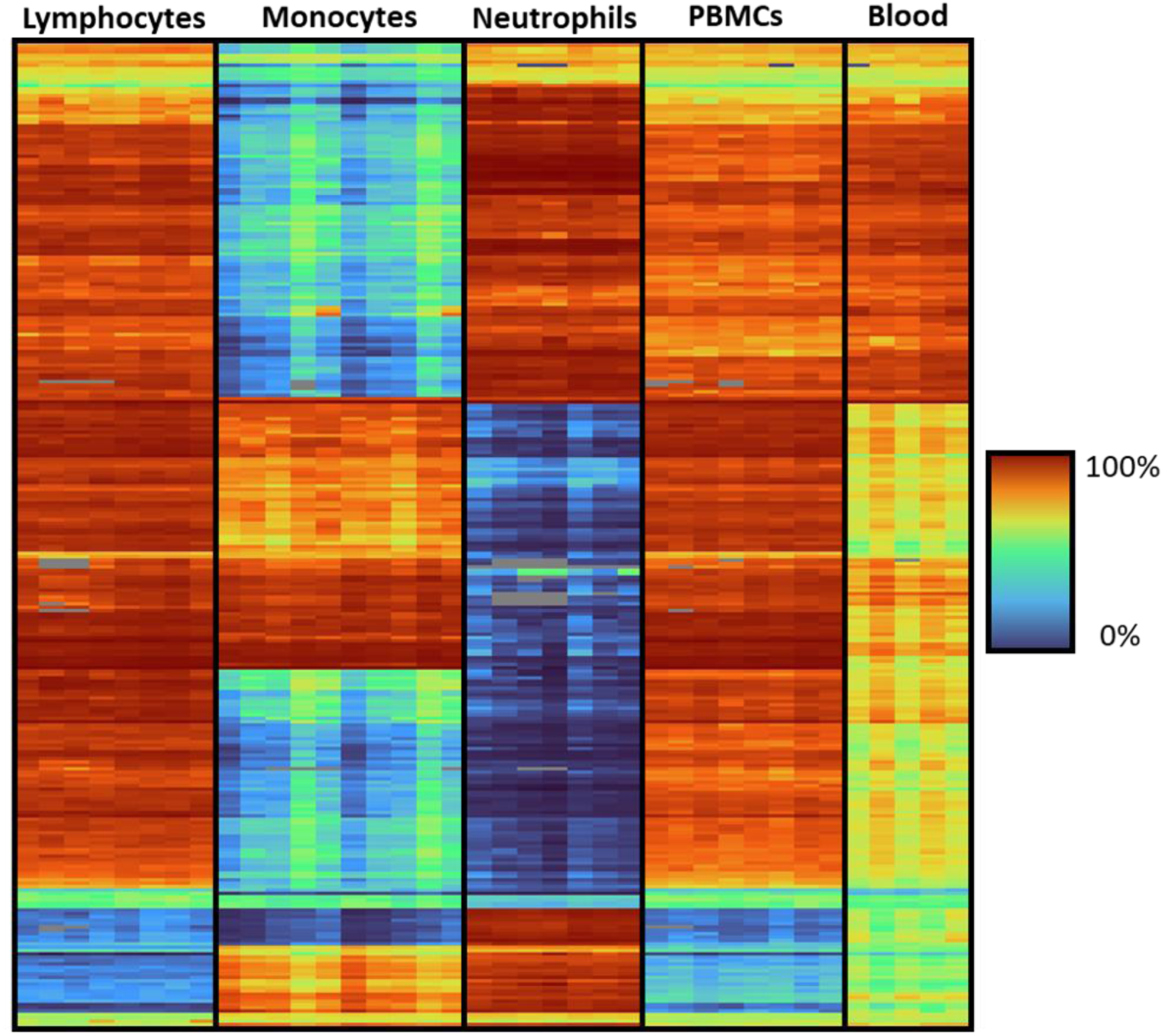
Cell-type specific DNA methylation patterns in blood. The heatmap represents DNA methylation values measured with the RUMIGEN EpiChip at 293 CpGs constituting the reference panel for deconvolution in purified lymphocytes (4 animals and two technical replicates per animal), monocytes (5 animals and two technical replicates per animal), neutrophils (4 animals and 1-2 technical replicates per animal), in PBMCs (4 animals and two technical replicates per animal) and whole blood (5 animals, no replicate).

**Figure 8:**
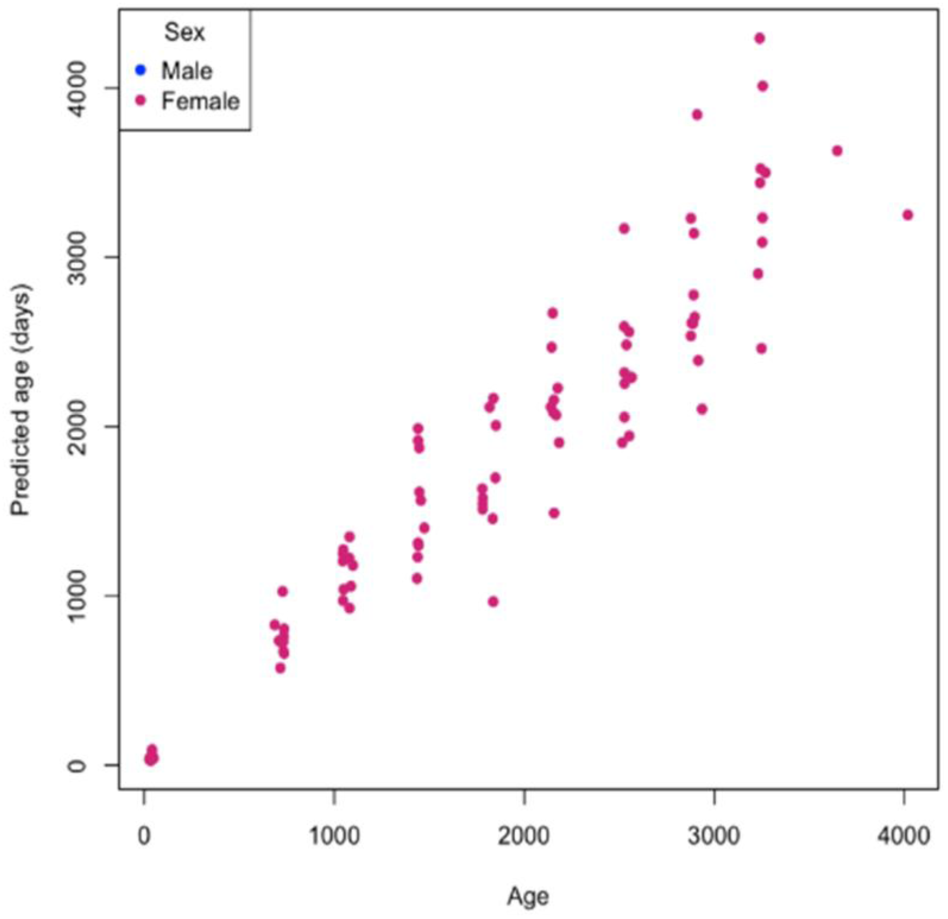
Epigenetic Clock. Chronological age (x-axis) versus epigenetic age estimated for clock model using log transformed chronological age.

### Epigenetic clock

A total of 40508 CpG sites were used for building a bovine epigenetic clock from the RUMIGEN EpiChip data obtained on 96 ear cartilage samples (41). Elastic net regression selected 205 CpG sites for the epigenetic clock. The predictive performance of the RUMIGEN EpiChip-based clock was highly accurate (R = 0.992) and a mean absolute error of 151 days with the accuracy of the estimators calculated using leave-one-out cross-validation.

## Discussion

### Design

The design of the EpiChip was guided by two main lines of reasoning. The first was to include CpG sites located within known regulatory genomic contexts, thereby enabling both fundamental studies of DNA methylation biology and analyses across tissues or cell types. This rationale is consistent with the design strategies used for the various versions of the human (13–15), and for the mouse DNA methylation BeadChips (42). These arrays feature dense coverage of gene bodies and promoters, and more recently, of distal intergenic regulatory elements. In our case, such depth of coverage was not feasible for two main reasons. First, the limited number of CpGs that could be incorporated within the probes constrained the achievable density of coverage. Second, at the time of design, the available genomic annotations were far less comprehensive than those for the human or mouse genomes.

Consequently, our design primarily focused on promoters, eQTLs and CTCF binding sites. In agreement with this strategy, the RUMIGEN EpiChip CpGs preferentially localize to promoters and TSS when compared to Ensembl standard annotation, reflecting the selection process during the design of the EpiChip. The signals of enrichment with novel annotation features were comparable to those obtained from randomly generated CpG sets, demonstrating that the EpiChip content is representative of the whole genome regarding these regulatory regions. Interestingly, several CpG sites co-localized with newly annotated regulatory regions detected across independent studies that spanned a diverse catalogue of tissue types (43–45). For example, we observed co-localization with bidirectional enhancer clusters annotated by the CAGE BovReg consortium (43), which encompass both tissue-specific and tissue-shared regulatory elements. Therefore, the RUMIGEN EpiChip CpGs preferentially target functionally relevant regulatory regions, effectively capturing epigenetic variation in both promoters and enhancers across multiple tissues, thereby underscoring their broad utility for functional epigenomic analyses.

The second design principle was to include CpG sites previously associated with phenotypic variation. This strategy was motivated by findings from Waterland’s group, who demonstrated that a large proportion of CpG sites included in the human EPIC array exhibit limited inter-individual variation within a given cell type (46). They hypothesized that this low variability reflects the initial selection of CpGs based on chromatin annotations—features that vary substantially across tissues and developmental stages within an individual, but not necessarily between individuals of the same tissue. This underscores the importance of including CpGs that are empirically known to vary among individuals, particularly in the context of EWAS. Accordingly, approximately 50% of the CpGs included on the EpiChip were selected based on their reported associations with various phenotypes and therefore exhibit measurable inter-individual variability. This enrichment for CpGs showing variable methylation across diverse phenotypes, environments and physiological situations is complementary to the design approach used for the pan-mammalian BeadChip, which targets 36K CpG sites located within highly conserved genomic regions across a wide range of mammalian species (47). This tool was specifically designed to allow inter-species comparisons, investigate evolutionary conserved epigenetic processes, and provide access to epigenetic profiling for species for which the development of a species-specific array would not be cost-effective for the numbers of samples that would potentially be analyzed.

### Technical performance

The design of the validation study was driven by the objective of assessing whether laboratory or treatment batches could influence DNA methylation beta-values. Batch effects in the context of microarray technologies are now well-documented in the literature, as the technology has been widely used over the past decades (48–50). To account for this, a substantial number of samples representing two cell types (semen and blood) were analyzed repeatedly on different positions on the same or different slides, across two experimental batches of 96 samples, and processed in two independent laboratories. A noteworthy feature of the two cell types used (semen and blood) is their extremely contrasted DNA methylation profiles, resulting from the extensive epigenetic reprogramming that occurs during germline differentiation (39, 40). This property allows our technical evaluation to cover a broad spectrum of the methylome using only two cell types. PCA revealed the presence of technical effects, both between laboratories and between batches within the same laboratory. Importantly, biological variation and technical variation were observed on orthogonal dimensions of the PCA, suggesting that they can easily be isolated from each other. In this study, a standard processing pipeline using the default parameters of the SeSaMe package was applied, which did not specifically correct for batch effects. Nevertheless, the literature offers a variety of methods developed to mitigate such effects (50–54). Although a detailed evaluation of correction methods was beyond the scope of the present work, they remain valuable tools for future analyses aimed at minimizing technical variation. Importantly, the presence of detectable batch effects did not substantially impact the repeatability or accuracy of the methylation measurements (beta-values), both of which were excellent. These findings align with previous assessments of similar technologies (16, 42). In addition, in both cell types, the EpiChip-derived beta-values showed strong concordance with those obtained using a gold standard sequencing-based technique, RRBS, for a subset of CpGs covered by both methods. This reinforced the reliability of DNA methylation measurements generated by EpiChip. It is worth noting that the high correlation between the two methods was largely driven by consistent agreement across the hypo- and hypermethylated sites (close to 0% and 100% DNA methylation, respectively), which is expected given the bimodal distribution of methylation in both cell types. Taken together, these results highlight the excellent technical performance of the RUMIGEN EpiChip and its capacity to measure biological variation with high reproducibility and accuracy, even in the presence of technical effects.

### Potential applications

The high precision of beta-value estimation offered by the RUMIGEN EpiChip and its relatively affordable cost, present a significant advantage in the context of EWAS studies, as it enables derivation of DNA methylation profiles for larger population samples, contributing to increase statistical power. Several studies have shown that, at specific loci, statistical power is often higher when using DNA methylation BeadChips compared to sequencing-based technologies (55, 56). This is primarily due to the fact that sequencing-based methylation estimates are highly dependent on coverage depth at each CpG site. Since WGBS or enzymatic methylation sequencing (EM-seq) remain costly, achieving high coverage across the genome is challenging. Even a coverage depth of 30X or more may still produce substantial technical noise, necessitating larger sample sizes to distinguish biological signal from background variability (56). In contrast, the EpiChip technology provides more consistent and precise methylation measurements at a much lower cost, enabling the analysis of larger cohorts. This combination of technical accuracy and cost-effectiveness makes this tool particularly well-suited for large-scale population studies and EWAS.

Moreover. the RUMIGEN EpiChip has also been designed to address a wide range of applications and biological questions, some examples are listed thereafter. Imprinted regions, particularly ICRs, are regions that regulate monoallelic gene expression depending on an epigenetic profile established during gametogenesis. Imprinted genes have been identified across numerous mammalian species including humans and model organisms, and have been associated with various developmental diseases, play critical roles in brain function, and are sensitive to environmentally induced epigenetic variation (57). As such, they may contribute to phenotypic variability in livestock and are of great interest to the epigenomics community (58). These regions are commonly included in existing DNA methylation microarrays such as the human EPIC BeadChip, the mouse methylation array, and more recently, the imprintome methylation BeadChip, which was developed to enhance coverage of imprinted loci (59). A growing share of bovine reproduction is being performed using embryo technologies, such as embryo transfer or *in vitro* fertilization, which are associated with developmental defects such as large offspring syndrome (60). Embryo technologies have been hypothesized to cause DNA methylation alterations at imprinted regions during early development, leading to placental dysfunction and growth abnormalities. The inclusion of imprinted regions in the EpiChip therefore offers the possibility to monitor the DNA status of these regions during normal and altered development.

Secondly, sets of probes were included on the EpiChip for their ability to discriminate between the three major blood cell types: neutrophils, lymphocytes, and monocytes (Costes et al., unpublished data), as well as between B cells, CD4 and CD8 T cells, γδ T cells, NK cells, monocytes and granulocytes (32). When combined with a reference methylation panel that provides DNA methylation values for each cell type at each locus, these markers could be used to perform deconvolution (61). This method infers the proportions of different cell types in a complex cell mixture, making it crucial for EWAS, as it is necessary to disentangled methylation variation due to cell composition from the one that is associated with the phenotype of interest. Furthermore, in cattle, routine access to detailed blood cell counts (blood formula) can be limited, so using DNA methylation as a proxy to estimate blood composition may offer a practical and informative approach, especially for studying associations with health traits, as it has been demonstrated in humans (62).

Finally, the EpiChip includes markers previously identified as components of a bovine epigenetic clock (41). The concept of the epigenetic clock is based on the observation that DNA methylation levels at specific CpG sites are strongly correlated with chronological age (63, 64). This metric has been associated with biological aging processes (65), and in the context of livestock production, may offer valuable insights into an animal’s physiological state. For instance, accelerated epigenetic aging could indicate poor adaptation to the environment or underlying health issues, thus providing useful guidance for animal management and breeding strategies.

### Perspectives on content evolution

One of the inherent limitations of array-based technologies is the finite number of CpG sites that can be interrogated simultaneously. However, this limitation could be partially overcome by the scalability of the technology. In humans, for example, successive updates of methylation arrays have demonstrated the feasibility of redesigning arrays based on newly available data and emerging scientific needs. The coming years will be crucial for generating large-scale datasets on various cell types using the current version of the RUMIGEN EpiChip. These data will provide valuable insights into the informativeness of individual CpG sites. Poorly performing probes and uninformative CpGs showing no DNA methylation variation can be removed from future designs. Eliminating such sites would free-up space to include novel probes targeting more variable or biologically relevant regions.

Another parameter for improving array performance involves optimizing the ratio between Type I and Type II probes. Indeed, Illumina recommended to maintain a proportion of Type I probes between 10% and 30% of the total content. However, Infinium I chemistry requires two probes in order to analyze one CpG site, compared to only one probe for Infinium II. Consequently, for a fixed array size of 50,000 probes, the number of CpG sites analyzed can vary from approximately 45,000 CpGs (with 10% Infinium I) to 42,500 CpGs (with 30% Infinium I). Determining the optimal balance between probe types could therefore help maximize CpG coverage while maintaining assay performance.

Finally, it is worth noting that, at the time of the EpiChip design, the bovine genome annotation was relatively limited compared to those of better-characterized species such as humans or mice. Since then, initiatives such as BovReg, CattleGTEx and FAANG, have significantly improved the functional annotation of the bovine genome (36, 43, 66). If additional space can be freed through probe optimization strategies, functionally important regions such as enhancers, insulators, or other regulatory elements identified by these projects could be enriched in an updated version of the array.

## Conclusion

We developed the RUMIGEN EpiChip, the first DNA methylation array specifically designed for cattle. The array includes 43,317 CpG sites, combining CpGs whose methylation levels vary with health status, physiological stage, fertility, and environmental challenges, along with CpGs located within functional genomic elements. This new resource demonstrates excellent technical performance in terms of repeatability, accuracy, and concordance with existing technologies. We also established its utility for investigating a range of biological processes in which DNA methylation plays a key role. Thanks to its compact format, the RUMIGEN EpiChip opens the way for routine applications similar to genotyping arrays, enabling integration of methylation data into management and breeding practices for both breeders and farmers.

## Methods

### CpG collection for the biomarker-based approach

A list of CpGs of interest was constituted firstly by implementing complementary genome-wide sequencing approaches or harnessing publicly available dataset, to which independent lists of CpGs of interest provided by various research groups were aggregated. Information regarding the source and CpG numbers in each dataset are listed in Supplementary Table 1.

Reduced representation bisulfite sequencing (RRBS) data was obtained as described (26, 67) and aligned on an *in silico* bisulfite-converted reference genome using Bismark (68). Only CpGs covered by ≥10 reads were retained. Oxford Nanopore Technology (ONT) was applied as described (69). Methylation marks were extracted using Nanopolish v0.13.2 (70) and aligned to the reference genome using minimap2 v2.17-r941 (71). Differential methylation analysis was performed as described (30) using DSS (72). In addition, publicly available WGBS data were collected from the accessions GSE175855 (73) GSE119263 (74), GSE142472 (75) and GSE121758 (76). Alignment on the *in silico* bisulfite-converted reference genome, deduplication, extraction of methylation sites were performed using Bismark (68). Only CpGs covered by ≥10 reads were retained, except for embryo samples from GSE121758 for which this threshold was lowered to 5 reads, in order to cover more CpGs.

The reference bovine genome used in all sequence alignments was the ARS-UCD1.2 assembly. Sites that potentially coincided with polymorphic markers listed in the 1000 Bull Genome Project (run 9) were filtered out. After merging all the different lists and filtering out CpGs overlapping putative SNPs, 250K unique CpGs remained.

### Design

All the biomarkers from this CpGs collection were submitted to probe design by Illumina. The probes containing SNPs in the 3’ last 15 nucleotides, targeting a CpG that directly falls into a SNP, or that mapped to several locations on the reference bisulfite-converted genome were discarded. Next, the probe selection strategy was conducted to have a proportional representation of the different origins of biomarkers within the final design. For the biomarkers identified by RRBS, only probes that showed an *in silico* quality score >0.6 for Infinium II probes and >0.4 for Infinium I probes, were retained. For the ONT biomarkers, the same score thresholds applied. To reduce further the list of ONT biomarkers, CpGs for which DNA methylation levels displayed a standard deviation <10% in blood; or located outside a 1000 bp range of eQTLs or a promoter were filtered out. The same criteria have been applied for the WGBS biomarkers.

The distances between all CpGs were calculated and in regions where a CpG has recursively another CpG within 10bp, only one CpG (the one having a probe with the highest *in silico* quality score) was retained to avoid redundancy.

In order to design probes targeting functional elements, all CpGs that fell into promoters of uncovered genes were submitted to probes design. The same probe quality filters as described above for the biomarkers were applied. Only one CpG per promoter was retained, and was selected based on the *in silico* probe score, the location relative to CTCF sites, eQTLs or the TSS.

Finally, this list was completed by probes used for routine quality controls. To this end, sample independent control probes monitoring the staining, extension, hybridization, target removal, negative control, and bisulfite conversion controls have also been designed.

Illumina manufactured the EpiChip with a conversion rate of 80%, leading to a total of 43,317CpGs physically present in the final design.

### Annotation with genes and genomic regulatory features

To investigate the enrichment of CpGs present on the EpiChip with respect to genes and regulatory features, we computed ORs to assess both enrichment and depletion across different genomic annotations. First, we analyzed enrichment in canonical gene features, including exons, introns, transcription start sites (TSS_100bp; ±100 bp from TSS), promoters (-2000/+100 bp from TSS), and transcription termination sites (TTS_100bp; ±100 bp from TTS). These features were extracted from the reference GTF file obtained from Ensembl (release 110; ftp://ftp.ensembl.org/pub/release-110/gtf/bos_taurus/Bos_taurus.ARS-UCD1.2.110.gtf.gz) and are collectively referred to as the common annotation database.

In addition, we also assessed enrichment using recently generated annotation catalogs (referred as novel annotation catalogs), including: (i) open chromatin regions (ATAC) identified by ATAC-seq across more than 60 bovine tissue types (44); (ii) transcription start sites (TSS_CAGE) and bidirectional enhancer clusters (Enhancer_CAGE) detected by CAGE-seq in over 20 tissue types (43); (iii) regulatory elements from the Ensembl Regulatory Build (release 113; ftp://ftp.ensembl.org/pub/release-113/regulation/bos_taurus/ARS-UCD1.3/annotation/Bos_taurus.ARS-UCD1.3.regulatory_features.v113.gff3.gz), including newly annotated promoters (Promoter_Ensembl), enhancers (Enhancer_Ensembl), and open chromatin regions (Open_Chromatin_regions_Ensembl) annotated in eight different tissues.

For each feature, observed ORs were calculated by comparing the EpiChip content to a randomized background set of the same size, based on the number of CpGs overlapping regulatory regions in the feature set. Randomized regions were generated using the regioneR package while accounting for bovine chromosome sizes as a reference. This randomization procedure was repeated 1,000 times to obtain the mean OR and the corresponding 95% confidence interval.

### Tissue collection and DNA samples

All study methods were implemented in accordance with EU guidelines and regulations (Directive 2010/63/UE).

The bull samples originated from bulls selected for artificial insemination and were provided by commercial companies located in France (SYNETICS, GEN’IATEST, UMOTEST and EVAJURA). For animals maintained in INRAE facilities, the experimental protocols were approved by national ethics committee under the reference APAFIS # 11503-2017091411167913 (11/02/2017).

Jugular blood samples were collected in EDTA-coated vacutainer tubes. Whole blood was diluted in two volumes of sterile Dulbecco′s Phosphate Buffered Saline (DPBS; pH 7) without Ca^2+^ and Mg^2+^, slowly transferred on top of Ficoll gradient (Ficoll Paque Plus, Sigma Aldrich) in a 2:3 volume ratio and centrifuged at 25°C for 30 min at 1000xg. After centrifugation, two major layers (plasma and Ficoll layers) could be observed. The interphase containing peripheral blood mononuclear cells (PBMCs) was collected, washed in PBS and centrifuged again. One part of the resulting pellet constituted the PBMC fraction. A second part was used to purify CD14+ monocytes using CD14+ beads following the manufacturer’s instructions (Miltenyi Biotec), as well as a CD14-negative cell fraction containing the total lymphocyte population. The Ficoll layer containing erythrocytes and neutrophils was mixed with one volume of DPBS and centrifuged for 10 minutes at 177xg. After two washes with DPBS, the cell pellet was suspended in 5 ml DPBS, and 25 ml sterile ultrapure water was added to induce a hypotonic shock and eliminate erythrocytes. After 30 second incubation, 10 ml 3.6% (w/v) NaCl was added and mixed by inversion. The neutrophil fraction was then recovered by centrifugation. All cell pellets were washed, resuspended in lysis buffer (10 mM Tris-HCl, pH 7.5, 2% SDS, 0.1 mM EDTA, 0.4 M NaCl), immediately frozen in liquid nitrogen and stored at −20 °C until use.

Bovine genomic DNA was purified from semen, total blood or different blood subfractions as described in Perrier et al. (39). For semen samples, 50 mM dithiothreitol (DTT) and 0.5 µg glycogen were added to the lysis buffer. DNA concentration was measured using a Qubit 2.0 Fluorometer (Invitrogen) and DNA integrity was checked on a 1% agarose gel. To obtain a wide range of targeted DNA methylation values in order to investigate the accuracy of methylation measurements, DNA samples from fresh semen and whole blood collected on the same individual were mixed in various proportions for two bulls. All the DNA samples used in this study as well as the number of technical replicates are presented in Table 1.

**Table 1.** DNA samples used for the technical validation of the EpiChip.

| Sample name | Sample type | Individual | Sex | Age (months) | Breed | Replicate number | Figures |
| --- | --- | --- | --- | --- | --- | --- | --- |
| Blood sample #1 | PBMCs | Bull #1 | Male | 17 | Holstein | 24 | 3, 5, 6, |
| Blood sample #2 | PBMCs | Bull #2 | Male | 17 | Holstein | 24 | 3, 5, 6 |
| Semen sample #1 | Semen | Bull #3 | Male | NA | Montbéliarde | 12 | 3, 5, 6 |
| Semen sample #2 | Semen | Bull #4 | Male | NA | Montbéliarde | 12 | 3, 5, 6 |
| Mix #1_0_100 | 0% semen, 100% blood | Bull #5 | Male | NA | Montbéliarde | 2 | 3, 4 |
| Mix #1_10_90 | 10% semen, 90% blood | Bull #5 | Male | NA | Montbéliarde | 2 | 4 |
| Mix #1_20_80 | 20% semen, 80% blood | Bull #5 | Male | NA | Montbéliarde | 2 | 4 |
| Mix #1_30_70 | 30% semen, 70% blood | Bull #5 | Male | NA | Montbéliarde | 2 | 4 |
| Mix #1_40_60 | 40% semen, 60% blood | Bull #5 | Male | NA | Montbéliarde | 2 | 4 |
| Mix #1_50_50 | 50% semen, 50% blood | Bull #5 | Male | NA | Montbéliarde | 2 | 4 |
| Mix #1_60_40 | 60% semen, 40% blood | Bull #5 | Male | NA | Montbéliarde | 2 | 4 |
| Mix #1_70_30 | 70% semen, 30% blood | Bull #5 | Male | NA | Montbéliarde | 2 | 4 |
| Mix #1_80_20 | 80% semen, 20% blood | Bull #5 | Male | NA | Montbéliarde | 2 | 4 |
| Mix #1_90_10 | 90% semen, 10% blood | Bull #5 | Male | NA | Montbéliarde | 2 | 4 |
| Mix #1_100_0 | 100% semen, 0% blood | Bull #5 | Male | NA | Montbéliarde | 2 | 3, 4 |
| Mix #2_0_100 | 0% semen, 100% blood | Bull #6 | Male | NA | Montbéliarde | 2 | 3, 4 |
| Mix #2_10_90 | 10% semen, 90% blood | Bull #6 | Male | NA | Montbéliarde | 2 | 4 |
| Mix #2_20_80 | 20% semen, 80% blood | Bull #6 | Male | NA | Montbéliarde | 2 | 4 |
| Mix #2_30_70 | 30% semen, 70% blood | Bull #6 | Male | NA | Montbéliarde | 2 | 4 |
| Mix #2_40_60 | 40% semen, 60% blood | Bull #6 | Male | NA | Montbéliarde | 2 | 4 |
| Mix #2_50_50 | 50% semen, 50% blood | Bull #6 | Male | NA | Montbéliarde | 2 | 4 |
| Mix #2_60_40 | 60% semen, 40% blood | Bull #6 | Male | NA | Montbéliarde | 2 | 4 |
| Mix #2_70_30 | 70% semen, 30% blood | Bull #6 | Male | NA | Montbéliarde | 2 | 4 |
| Mix #2_80_20 | 80% semen, 20% blood | Bull #6 | Male | NA | Montbéliarde | 2 | 4 |
| Mix #2_90_10 | 90% semen, 10% blood | Bull #6 | Male | NA | Montbéliarde | 2 | 4 |
| Mix #2_100_0 | 100% semen, 0% blood | Bull #6 | Male | NA | Montbéliarde | 2 | 3, 4 |
| Whole blood #1 | Whole blood | Cow #1 | Female | 14 | Holstein | 1 | 7 |
| Whole blood #2 | Whole blood | Cow #2 | Female | NA | Holstein | 1 | 7 |
| Whole blood #3 | Whole blood | Cow #3 | Female | NA | Holstein | 1 | 7 |
| Whole blood #4 | Whole blood | Cow #4 | Female | NA | Holstein | 1 | 7 |
| Whole blood #5 | Whole blood | Cow #5 | Female | NA | Holstein | 1 | 7 |
| PBMC #1 | PBMCs | Cow #6 | Female | 80 | Holstein | 2 | 7 |
| PBMC #2 | PBMCs | Cow #7 | Female | 78 | Holstein | 2 | 7 |
| PBMC #3 | PBMCs | Cow #8 | Female | 74 | Holstein | 2 | 7 |
| PBMC #4 | PBMCs | Cow #9 | Female | 74 | Holstein | 2 | 7 |
| Monocyte #1 | CD14+ | Cow #10 | Female | NA | Holstein | 2 | 7 |
| Monocyte #2 | CD14+ | Cow #11 | Female | NA | Holstein | 2 | 7 |
| Monocyte #3 | CD14+ | Cow #12 | Female | NA | Holstein | 2 | 7 |
| Monocyte #4 | CD14+ | Cow #13 | Female | 81 | Holstein | 2 | 7 |
| Monocyte #5 | CD14+ | Cow #14 | Female | 79 | Holstein | 2 | 7 |
| Lymphocyte #1 | CD14 minus | Cow #15 | Female | 81 | Holstein | 2 | 7 |
| Lymphocyte #2 | CD14 minus | Cow #16 | Female | 81 | Holstein | 2 | 7 |
| Lymphocyte #3 | CD14 minus | Cow #17 | Female | 78 | Holstein | 2 | 7 |
| Lymphocyte #4 | CD14 minus | Cow #9 | Female | 74 | Holstein | 2 | 7 |
| Neutrophil #1 | Neutrophils | Cow #18 | Female | NA | Holstein | 2 | 7 |
| Neutrophil #2 | Neutrophils | Cow #19 | Female | NA | Holstein | 2 | 7 |
| Neutrophil #3 | Neutrophils | Cow #20 | Female | NA | Holstein | 2 | 7 |
| Neutrophil #4 | Neutrophils | Cow #16 | Female | 81 | Holstein | 2 | 7 |

NA, information not available. PBMCs, peripheral blood mononuclear cells

### Bisulfite conversion and array hybridization

Each DNA sample was prepared to reach a concentration of 50 ng/µl.

Bisulfite conversion was carried out by GD Biotech genotyping facility (France) using the Zymo EZ DNA Methylation-Lightning Automation Kit. The first steps were carried out manually: 1 µg of genomic DNA was mixed with the Lightning Conversion Reagent, and the conversion reaction was performed in a thermocycler (Labcycler Gradient, SensoQuest) according to the manufacturer’s recommended thermal profile. Then, the automated-elution run was conducted using the MagnetaPure96 system, following the manufacturer’s protocol. The converted DNA was eluted in 25 µL elution buffer and stored at –20°C.

Epigenotyping was performed using the Infinium HTS Methylation Assay kit, in accordance with the manufacturer’s instructions. 4 µL of bisulfite-converted DNA was first denatured under alkaline conditions, neutralized and then amplified by isothermal whole-genome amplification (iWGA), which was carried out between 20 and 24h at 37 °C in a temperature-controlled Illumina oven. After amplification, DNA was enzymatically fragmented at 37 °C, precipitated at 4 °C with 100% 2-propanol and dried at room temperature. The DNA pellet was resuspended in hybridization buffer and loaded onto the arrays after denaturation at 95 °C. Hybridization was performed at 48 °C for 16 hours in a temperature-controlled Illumina oven, in accordance with the kit specifications. Single base extension, allowing the addition of a single labeled base to each probe, and staining, involving antibody binding and red/green fluorescent labeling to discriminate between methylated/unmethylated status and amplify the signal, was executed automatically using the Tecan Freedom EVO robotic workstation. Fluorescent signal detection was performed using the iScan System (Illumina), following the manufacturer’s guidelines for image acquisition and data quality control.

A subset of samples (blood samples #1 and #2, semen samples #1 and #2 and semen-blood mixes) was treated by a second genotyping facility, Xenética Fontao (Spain), from 250 ng DNA using the Zymo EZ-96 DNA Methylation-Lightning™ MagPrep kit. The protocol used for processing the bisulfite-converted DNA and for array hybridization was then the same as described above.

### Data processing and quality controls

Raw intensity data (.idat files) were preprocessed using the R package SeSAME (77) and the function openSeSaMe() with “prep” argument assigned to “QCDPB”. This step allowed the normalization of the data, and included the suppression of probes of poor design (Q), the inference of Infinium channel (C), dye bias correction (D), the detection of probes of poor quality by pOOBAH and the background correction using noob (B). Finally, preprocessed signals were translated into beta values. After conversion, probe signals that were assimilated to background by pOOBAH were set as “NA” in the corresponding sample. Probes displaying “NA” in more than 20% of the analyzed samples were then removed from subsequent analysis.

For quality controls, the fluorescence data was analyzed using readIDATpair() function prior to normalization. For the sample independent controls (“staining”, “extension”, “hybridization”, “target removal”), fluorescent data was consistent with the expected values (GenomeStudio documentation, https://www.illumina.com/content/dam/illumina-support/documents/documentation/software_documentation/genomestudio/genomestudio-2011-1/genomestudio-methylation-v1-8-user-guide-11319130-b.pdf). Negative controls showed a minimal fluorescence signal in every channel as expected. Bisulfite controls displayed a high intensity in the Unmethylated probe channel (U) and a minimal intensity in the Methylated probe channel (M), demonstrating a correct bisulfite conversion.

### Epigenetic clock

For the epigenetic clock construction, ear punch samples (Allflex tissue sampling units (TSU)) and subsequent DNA extractions from a total of 96 cows aged 1 month to 11 years and month old from the Livestock Improvement Corporation (LIC) Ltd dairy farm was carried out as previously described by Caulton et.al. (41). Bisulfite conversion of 1ug input DNA was carried out with EZ DNA MethylationGold Kit (Zymo Research) following the manufacturer’s instructions. Array hybridisation and staining were performed with an automated liquid handling robot (Tecan Trading AG) and array scanning and imaging were executed on the Illumina Iscan platform at GenomNZ, AgResearch, NZ.

CpG sites were filtered to retain only those with valid beta values in at least 80% of samples, whereas samples were filtered out if greater that 10% milling values. Missing beta values were imputed using the mean value for the CpG site. Epigenetic clock construction was based on epigenetic “Clock 1” by Caulton et.al. (41). Briefly, the R package glmnet (78) within R (version 4.5.1) was used to implement an elastic-net regression model (α=0.5, γ = lambda.min) that included log-transformation of chronological age but no age offset as all animals in the study were sampled after birth. The accuracy of the estimators was calculated using leave-one-out cross-validation (LOOCV) by executing the cv.glmnet() program on each set of n − 1 samples, where n is the number of samples. The predicted age of the omitted sample was calculated with the model built using data from the remaining samples.

## Abbreviations

WGBS: whole genome bisulfite sequencing;
SNP: single nucleotide polymorphism;
RRBS: reduced representation bisulfite sequencing;
EWAS: epigenome-wide association studies;
ONT: Oxford Nanopore Technology;
TSS: transcription start site;
eQTL: expression quantitative trait locus;
CTCF: CCCTC-binding factor;
OR: odds ratio;
TTS: transcription termination site;
CAGE: cap analysis of gene expression;
ATAC: assay for transposase-accessible chromatin;
PCA: principal component analysis;
ICR: imprinting control region;
EM-seq: enzymatic methylation sequencing.

## Ethics approval and consent to participate

All study methods were implemented in accordance with EU guidelines and regulations (Directive 2010/63/UE). The bull samples originated from bulls selected for artificial insemination and were provided by commercial companies. For animals maintained in INRAE facilities, the experimental protocols were approved by national ethics committee under the reference APAFIS #11503-2017091411167913 (11/02/2017).

## Consent for publication

Not applicable

## Availability of data and materials

All data supporting the findings of this study are available within the paper and its Supplementary Information. The word file Supplementary information_Costes_Epichip.docx includes Supplementary Table 1 (CpG collection submitted to probe design for the biomarker-based strategy), Supplementary Figures 1 to 4 and Supplementary Table 4 (Gene imprinted regions and CpG coordinates used in Figure 6). The text files Supplementary_Table2.txt and Supplementary_Table3.txt list the CpG positions included on the EpiChip, together with their coordinates on the cattle genome (ARS-UCD1.2 assembly) and annotation features. The text file Supplementary_Table5.txt lists the CpG markers (probe IDs) together with the beta-values for each sample. The text file Supplementary_Table6.txt describes the metadata associated to samples in Supplementary_Table5.txt. The text file Supplementary_Table7.txt provides the genomic coordinates associated to probe IDs for all CpG markers listed in Supplementary_Table5.txt. RRBS fastq files used for the comparison with the RUMIGEN EpiChip are available through the European Nucleotide Archive (ENA) at EMBL-EBI under accession number PRJEB105696.

## Authors’ contributions

VC performed the design of the RUMIGEN EpiChip, performed data analysis and drafted the manuscript. ALC performed experiments and data analysis and contributed to the RUMIGEN EpiChip content. ARRS performed experiments and data analysis. GCMM annotated the EpiChip. SM, LL, CGA, JLV, ACT, FA performed experiments. CC performed data analysis. MB, SF, EMI-A, NB, JP, PN, AC, KT, EK contributed to the RUMIGEN EpiChip content. GF and DR provided samples. CLD obtained funding and provided samples. LS obtained funding and designed the study. ZC, GS, SC, HJ and OGR provided samples and contributed to the RUMIGEN EpiChip content. CP obtained funding and coordinated the study. HK obtained funding, coordinated the study, provided samples, contributed to the RUMIGEN EpiChip design and content, and drafted the manuscript. All authors revised the manuscript and approved the submitted version.

## Acknowledgement

We thank the breeding companies and experimental farms that provided biological samples and Illumina for the probe design and the technical support during implementation of epigenotyping. We are grateful to the Genotoul bioinformatics platform Toulouse Occitanie (Bioinfo Genotoul, https://doi.org/10.15454/1.5572369328961167E12) for providing computing and storage resources.

