## Supplementary_data for "The RUMIGEN EpiChip: a versatile, medium density DNA methylation Beadchip for large scale population studies in cattle"

| Cell type | Study | Number of samples | Sex | Age/stage | Breed | DNA methylation analysis | Number of CpGs | Reference |
|---|---|---|---|---|---|---|---|---|
| Monocytes | Age | 8 | Female | 6 years vs. 10-15 years | Holstein | RRBS | 300 | |
| Monocytes, lymphocytes, neutrophils | Immune cell types (deconvolution) | 36 | Female | 6-10 years | Holstein | RRBS | 400 | |
| B cells, CD4 αβ T cells, CD8 αβ T cells, γδ T cells, NK cells, monocytes and granulocytes | Immune cell types (deconvolution) | 59 | Female | 10-141 months | Holstein Friesian, N'Dama, Nelore | RRBS | 647 | (1) |
| Monocytes | Inflammatory status (LPS treatment) | 32 | Female | 6 years | Holstein | RRBS | 470 | |
| Monocytes | Physiological stage (15 days post-estrus after hormonal stimulation and ovulation vs. 15 days (±2) post calving) | 16 | Female | 6 years | Holstein | RRBS | 264 | |
| Monocytes | Nutrition | 6 | Female | NA | Holstein | RRBS | 542 | |
| Whole blood | Heat stress | 6 | Female | NA | Holstein | ONT | 7113 | |
| PBMCs | Mastitis sensitivity | 6 | Female | 44-54 months | Holstein | ONT | 8482 | |
| Whole blood | Metabolic stress | 6 | Female | 0-15 days | Holstein | ONT | 72982 | |
| Whole blood | Breed | 4 | Female | NA | Jersey, Holstein | ONT | 18633 | |
| Whole blood | Metabolic stress | 8 | Female | 0-2 days | NA | WGBS | 291 | GSE175855 (2) |
| Semen | Age | 18 | Male | 11-31 vs.50-83 months | Holstein | RRBS | 2058 | |
| Semen | Breed | 35 | Male | 11-83 months | Holstein, Montbéliarde, Normande, Charolaise, Abondance | RRBS | 9177 | |
| Semen | Early life nutrition | 25 | Male | 15-16 months | Holstein | RRBS | 303 | (3) |
| Semen | Fertility | 138 | Male | 18 months | Montbéliarde, Holstein | RRBS | 5697 | (4) |
| Semen | Fertility | 20 | Male | 18-145 months | Holstein | RRBS | 234 | (5) |
| Semen | Fertility | 11 | Male | 17-40 months | Belgian Blue | RRBS | 1744 | |
| Semen | Fertility | 20 | Male | 16-20 months | Abondance | RRBS | 199 | |
| Semen | Fertility | 8 | Male | 16-19 months | Tarentaise | RRBS | 3107 | |
| Semen | Season/age | 20 | Male | 20, 26, 32, 39, 49 months | Belgian Blue | RRBS | 1980 | |
| Semen | Fertility | 12 | Male | 10-42 months | Holstein, Jersey | ONT | 10234 | (6) |
| Semen | Breed | 12 | Male | 10-42 months | Holstein, Jersey | ONT | 1005 | |
| Semen | Fertility | 17 | Male | 10-144 months | Japanese Black | EPIC | 19 | (7) |
| Semen | Fertility | 6 | Male | 11-14 months | Holstein | WGBS | 186 | GSE119263 (8) |
| Semen | Fertility | 10 | Male | NA | NA | WGBS | 226 | GSE142472 (9) |
| Blastocysts | Post-fertilization eeprogramming | 12 | NA | 6-7 days after IVF | Holstein | RRBS | 15328 | |
| Embryos | Post-fertilization reprogramming | 16 | NA | 2–4 days after estrus (2 cells, 4 cells, 8 cells, 16 cells embryos) | Holstein | WGBS | 304 | GSE121758 (10) |
| Ileum, ileal lymph node | Gut immunity (MAP infection) | 7 | Female | NA | Holstein | WGBS | 27528 | (11) |
| Mammary gland | Milk quality | 17 | Female | NA | Holstein | WGBS | 1145 | (12) |
| Milk somatic cells | *Streptococcus uberis* subclinical mastitis | 6 | Female | NA | Holstein | WGBS | 10170 | (13) |

| | | | | | | | | |
|---|---|---|---|---|---|---|---|---|
| Milk somatic cells | *Staphylococcus aureus* subclinical mastitis | 26 | Female | 21-80 months | Holstein | WGBS | 38893 | (14) |
| Milk somatic cells | *Staphylococcus chromogenes* subclinical mastitis | 8 | Female | NA | Holstein | WGBS | 7258 | (15) |
| Ear cartilage | Aging (epigenetic clock) | 96 | Female | 1 month to 11 years and 1 month | Kiwicross | HorvathMammalMethyl40 | 441 | (16) |

**Supplementary Table 1.** CpG collection submitted to probe design for the biomarker-based strategy. The CpG numbers indicated for each study was obtained after filtering out overlapping SNPs, except for deconvolution (1), epigenetic clock (16) and Japanese black (7) studies. For two studies having identified a large number of candidate biomarkers using WGBS (14, 15), only CpGs included in promoters/first exon were considered, with the aim of narrowing down the list while maximizing the likelihood of capturing CpGs involved in gene regulation. Some CpGs were identified in different comparisons, but the integration of all the studies led to a collection of unique 250K CpGs. PBMCs, peripheral blood mononuclear cells; LPS, lipopolysaccharide; RRBS, reduced representation bisulfite sequencing; ONT, Oxford Nanopore Technology; WGBS, whole genome bisulfite sequencing; MAP, *Mycobacterium avium subsp. paratuberculosis* infection.

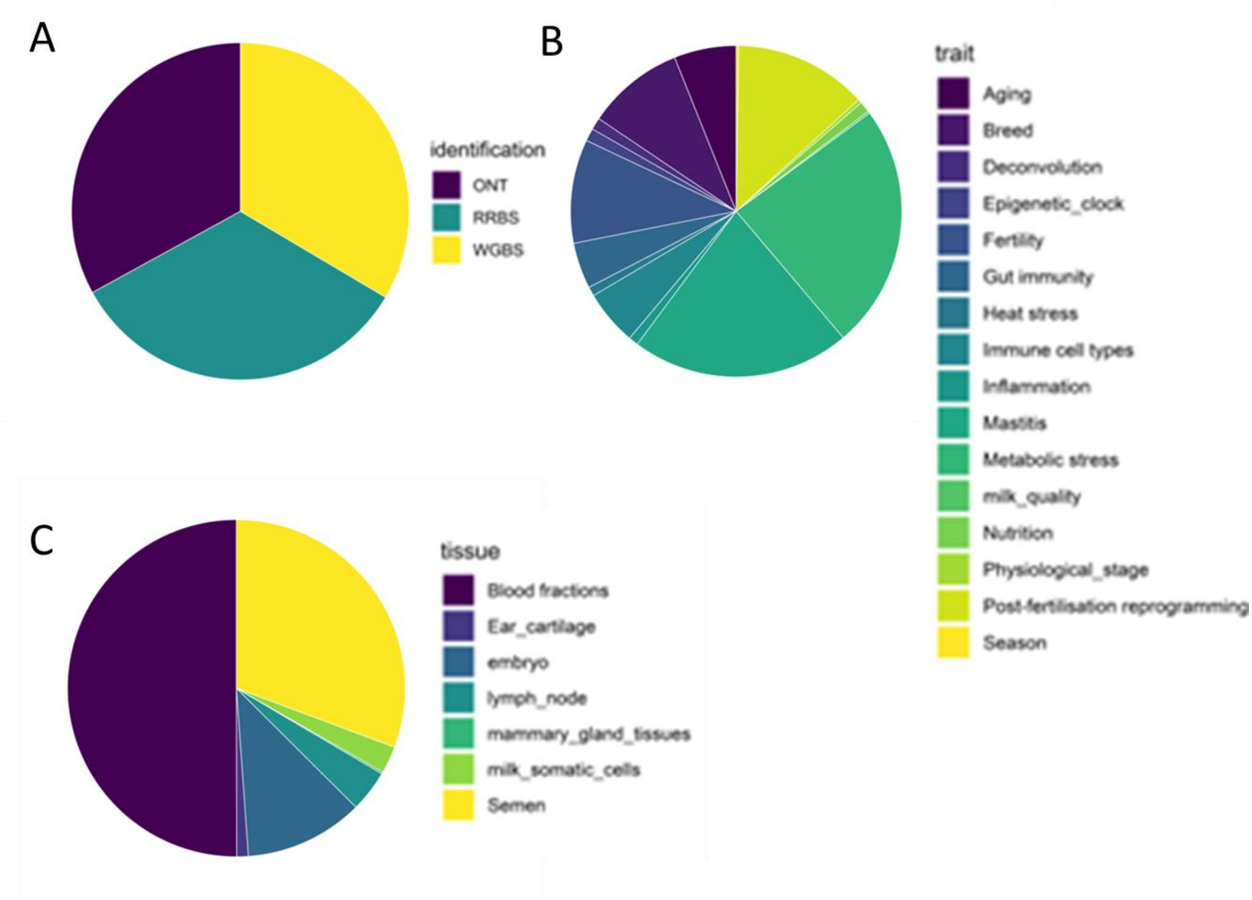


**Supplementary Figure 1.** Pie chart showing the origin of the selected biomarkers, indicating the technologies by which they were identified (A), the studies in which they were highlighted (B), and the tissues in which they were detected (C).

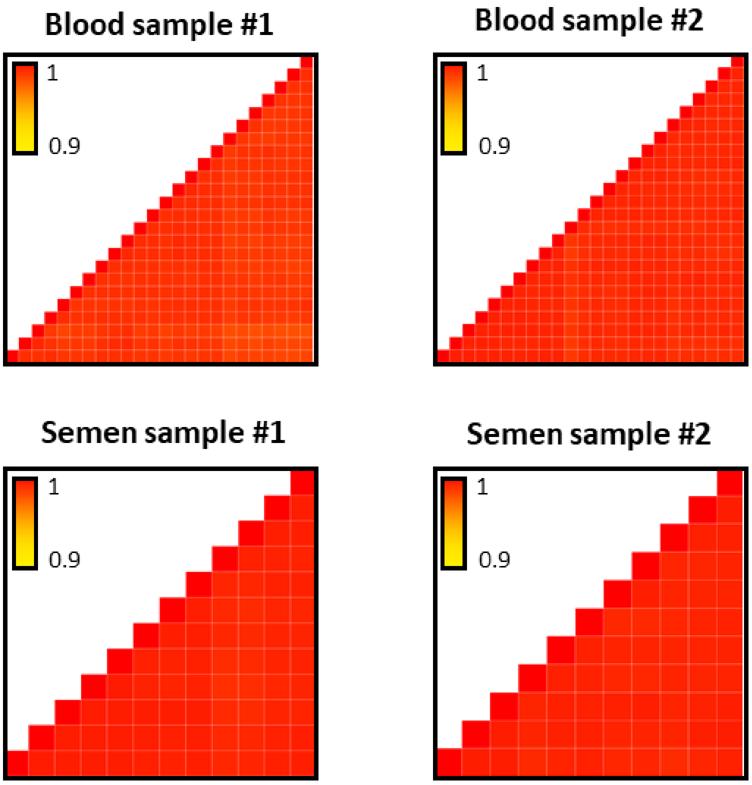


**Supplementary Figure 2.** Repeatability of DNA methylation measurements across technical replicates. Two blood (blood samples #1 and #2) and semen (semen samples #1 and #2) samples were analyzed using the RUMIGEN EpiChip with respectively 24 technical replicates for each PBMC sample and 12 technical replicates for each semen sample. For each of the four samples the Pearson correlation coefficients among technical replicates was calculated. The results are presented as a correlation matrix, where replicates are in the same order in rows and columns and where the color of each square represents the value of the correlation coefficient among two replicates. The mean correlation was 0.993 for blood sample #1, 0.996 for blood sample #2 and 0.997 for both semen sample #1 and semen sample #2.

| **Gene** | **CpG numbers** | **CpG coordinates** | **References** |
|---|---|---|---|
| *GNAS* | 5 | Chr13:57530764-57530770-57530775-57530891-57530893 | (17) |
| *GNAS* promoter | 5 | Chr13:57486529-57486550-57486808-57486844-57486120 | (18) |
| ICR *H19/IGF2* | 1 | Chr29: 49505364 | (19) |
| *IGF2R* | 3 | Chr9:96222522-96222172-96222386 | (20) |
| *PEG10* | 2 | Chr4:12063959-12064556 | (20) |
| *PEG3* | 14 | Chr18:64121303-64121323-64121359-64121445-64121461-64120458-64120470-64120482-64121308-64121347-64121362-64121447-64121455-64122290 | (21) |
| *PLAGL1* | 5 | Chr9:81419234-81419315-81419343-81419284-81419303 | (20) |
| *SNRPN* | 2 | Chr21:1938001-1938741 | (20) |

**Supplementary Table 4.** Gene imprinted regions and CpG coordinates used in Figure 6.

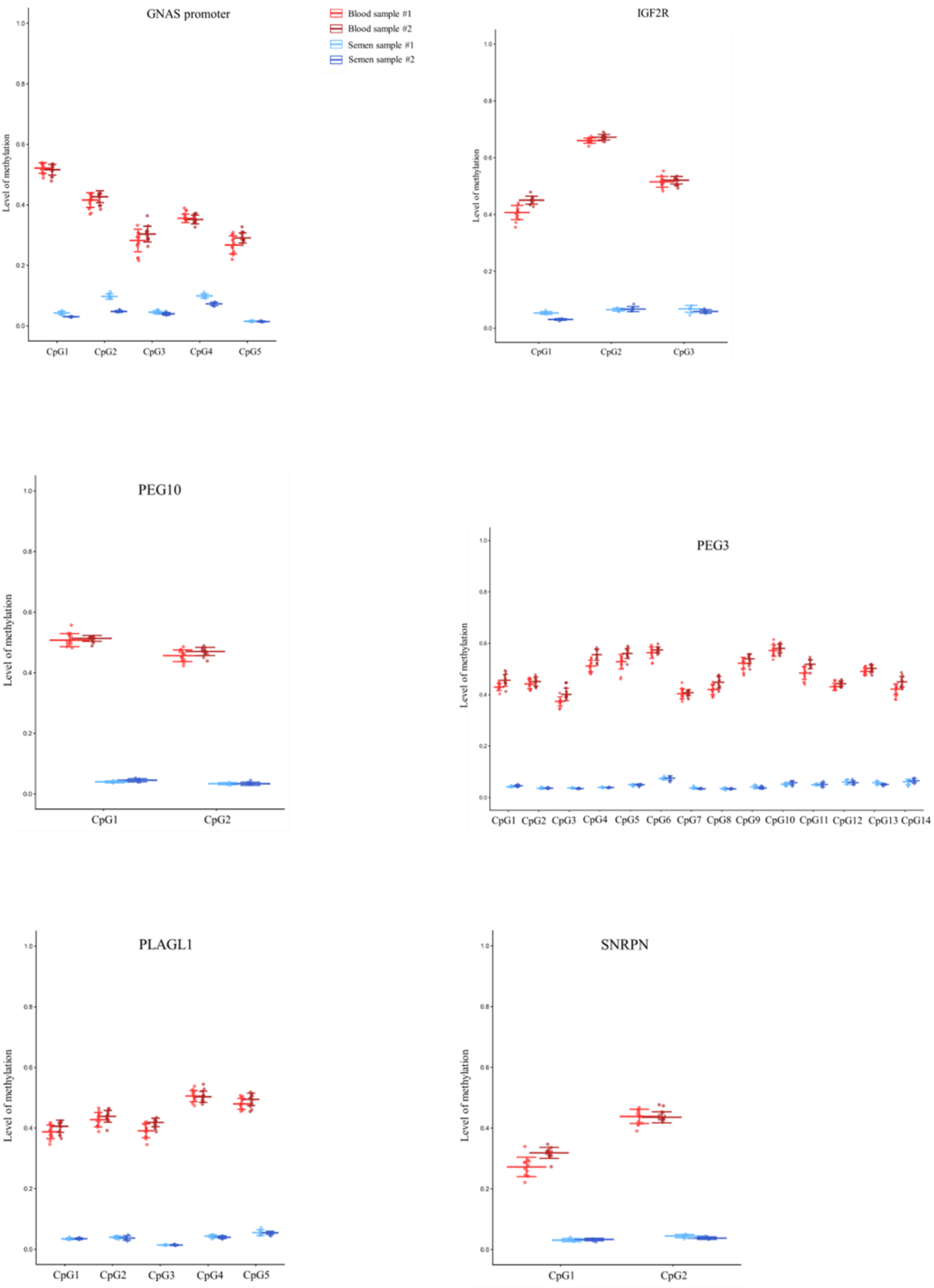


**Supplementary Figure 3.** DNA methylation at imprinted regions in 6 genes analyzed using the RUMIGEN EpiChip. Technical replicates of four independent samples (blood samples #1 and #2 and semen samples #1 and #2, Table 2) from two different cell types (semen and blood) and two animals, were analyzed by one laboratory in one single batch. DNA methylation values are displayed at the level of single CpGs, revealing heterogeneity across CpGs but homogeneity within biological and technical replicates.

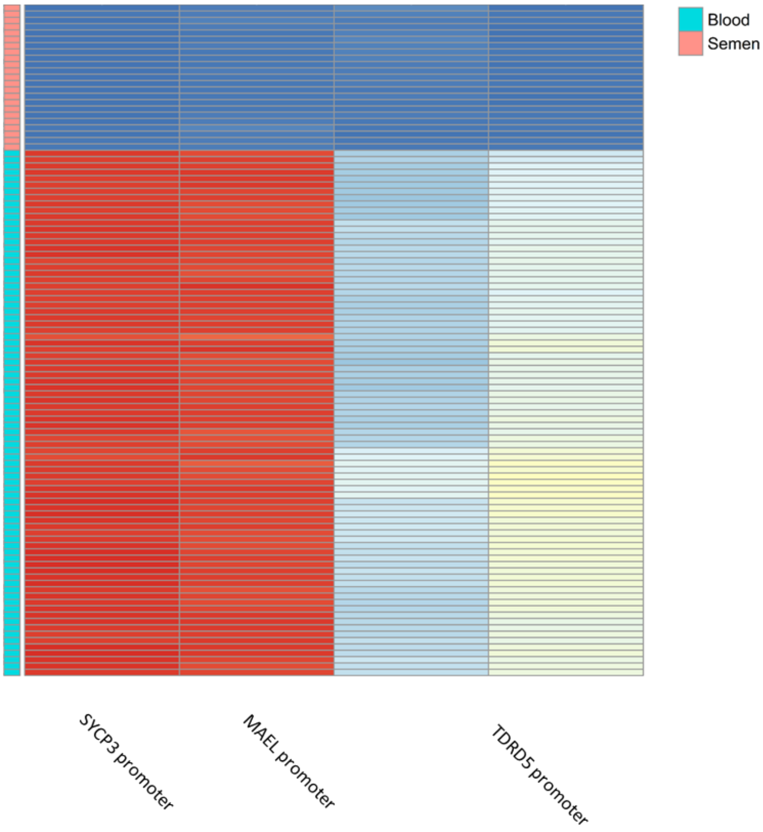


**Supplementary Figure 4.** Heatmap representing DNA methylation values measured with the RUMIGEN EpiChip at 4 CpGs located in promoters of three germinal genes, in blood and semen samples. The semen samples are consistently less methylated than the blood samples at all CpGs analyzed.

1. Powell J, Talenti A, Fisch A, Hemmink JD, Paxton E, Toye P, et al. Profiling the immune epigenome across global cattle breeds. Genome biology. 2023;24(1):127.
2. Zhang Y, Chaput C, Fournier E, Prunier J, Sirard MA. Comparing the whole genome methylation landscape of dairy calf blood cells revealed intergenerational inheritance of the maternal metabolism. Epigenetics. 2022;17(7):705-14.
3. Perrier JP, Kenny DA, Chaulot-Talmon A, Byrne CJ, Sellem E, Jouneau L, et al. Accelerating Onset of Puberty Through Modification of Early Life Nutrition Induces Modest but Persistent Changes in Bull Sperm DNA Methylation Profiles Post-puberty. Frontiers in genetics. 2020;11:945.
4. Costes V, Chaulot-Talmon A, Sellem E, Perrier JP, Aubert-Frambourg A, Jouneau L, et al. Predicting male fertility from the sperm methylome: application to 120 bulls with hundreds of artificial insemination records. Clinical epigenetics. 2022;14(1):54.
5. Stiavnicka M, Chaulot-Talmon A, Perrier JP, Hosek P, Kenny DA, Lonergan P, et al. Sperm DNA methylation patterns at discrete CpGs and genes involved in embryonic development are related to bull fertility. BMC genomics. 2022;23(1):379.
6. López-Catalina A, Costes V, Peiró-Pastor R, Kiefer H, González-Recio O. Oxford nanopore sequencing as an alternative to reduced representation bisulphite sequencing for the identification of CpGs of interest in livestock populations. Livestock Science. 2024;279.
7. Takeda K, Kobayashi E, Ogata K, Imai A, Sato S, Adachi H, et al. Differentially methylated CpG sites related to fertility in Japanese Black bull spermatozoa: epigenetic biomarker candidates to predict sire conception rate. The Journal of reproduction and development. 2021;67(2):99-107.
